# Genetic diversity, determinants, and dissemination of *Burkholderia pseudomallei* lineages implicated in melioidosis in northeast Thailand

**DOI:** 10.1101/2023.06.02.543359

**Authors:** Rathanin Seng, Chalita Chomkatekaew, Sarunporn Tandhavanant, Natnaree Saiprom, Rungnapa Phunpang, Janjira Thaipadungpanit, Elizabeth M Batty, Nicholas PJ Day, Wasun Chantratita, T. Eoin West, Nicholas R Thomson, Julian Parkhill, Claire Chewapreecha, Narisara Chantratita

## Abstract

Melioidosis is an often-fatal neglected tropical disease caused by an environmental bacterium *Burkholderia pseudomallei*. However, our understanding of the disease-causing bacterial lineages, their dissemination, and adaptive mechanisms remains limited. To address this, we conducted a comprehensive genomic analysis of 1,391 *B. pseudomallei* isolates collected from nine hospitals in northeast Thailand between 2015 and 2018, and contemporaneous isolates from neighbouring countries, representing the most densely sampled collection to date. Our study identified three dominant lineages with unique gene sets enhancing bacterial fitness, indicating lineage-specific adaptation strategies. Crucially, recombination was found to drive lineage-specific gene flow. Transcriptome analyses of representative clinical isolates from each dominant lineage revealed heightened expression of lineage-specific genes in environmental versus infection conditions, notably under nutrient depletion, highlighting environmental persistence as a key factor in the success of dominant lineages. The study also revealed the role of environmental factors – slope of terrain, altitude, direction of rivers, and the northeast monsoons – in shaping *B. pseudomallei* geographical dispersal. Collectively, our findings highlight persistence in the environment as a pivotal element facilitating *B. pseudomallei* spread, and as a prelude to exposure and infection, thereby providing useful insights for informing melioidosis prevention and control strategies.

## Introduction

Melioidosis, a severe infectious disease, affects an estimated 165,000 cases globally each year, of which 89,000 are fatal^1^. The disease is caused by *Burkholderia pseudomallei*, a Gram-negative bacillus found in soil and contaminated water across tropical and sub-tropical regions. Historically, limited access to microbiology laboratories for culture-confirmed diagnosis led to underreporting, particularly in lower- and middle-income countries^2^. However, improved infrastructure and awareness have led to increases in reported cases across South Asia, Southeast Asia, East Asia^3–8^ and Australia^9,10^ In Southeast Asia, the disease incidence is often linked to agriculture practice, particularly during the rainy seasons when rice paddy fields are flooded for planting. The flooded terrain enables the bacterium in the soil to surface, potentially exposing farmers to *B. pseudomallei* and subsequently leading to melioidosis^11^. Additionally, many cases of melioidosis have been associated with severe weather events^12–14^. While climate likely influences human encounters with *B. pseudomallei*, further investigation is needed to fully understand the mechanisms linking environmental factors to melioidosis epidemiology.

Understanding the population structure, dissemination and adaptation of *B. pseudomallei* in these climatically-challenged endemic regions requires a large-scale, geographically and chronologically densely-sampled, genetic dataset. Previous studies, albeit limited in sample size, have demonstrated that *B. pseudomallei* dissemination is driven by both anthropogenic and environmental factors^15–18^. Streams^19,20^, monsoons, typhoons and cyclones^13,14,21,22^ were identified as significant contributors to bacterial dissemination, highlighting the importance of bacterial persistence across a range of environmental conditions. *B. pseudomallei* exhibits remarkable survival capabilities across diverse environments, spanning from wet to dry, nutrient-depleted soil^23–28^ thereby enabling the bacterium to thrive in various ecological niches. Previous studies have noted the temporal and geographical co-existence of multiple *B. pseudomallei* lineages^15,16,29^. However, little is known about their distinct genetic content and adaptive strategies. Identification of lineage-specific genes associated with bacterial persistence and disease escalation will be essential to develop disease control strategies.

In this study, we conducted a population genomics analysis using combined *B. pseudomallei* isolates from melioidosis patients across nine provinces in the northeast Thailand including Buriram, Khon Kaen, Mahasarakam, Mukdahan, Nakhon Phanom, Roi-Et, Sisaket, Surin and Udon Thani^8^; totaling 1,265 isolates collected from July 2015 to December 2018. Additionally, we incorporated contemporary environmental and clinical collections from Thailand and neighbouring countries^15,29–34^, consisting of 15 clinical isolates and 111 environmental isolates (**Figure 1a**, **Supplementary data 1**). Our comprehensive analysis, including a total of 1,391 isolates, revealed the population structure, dissemination patterns, and genetic diversity of this bacterium. We identified genetic determinants associated with dominant lineages and investigated their biological functions and expression conditions in three representative isolates, each representing a dominant lineage. This provides insights into the strategies employed by *B. pseudomallei* lineages for successful persistence in the environment, ultimately leading to human exposure and infection.

**Figure 1.**
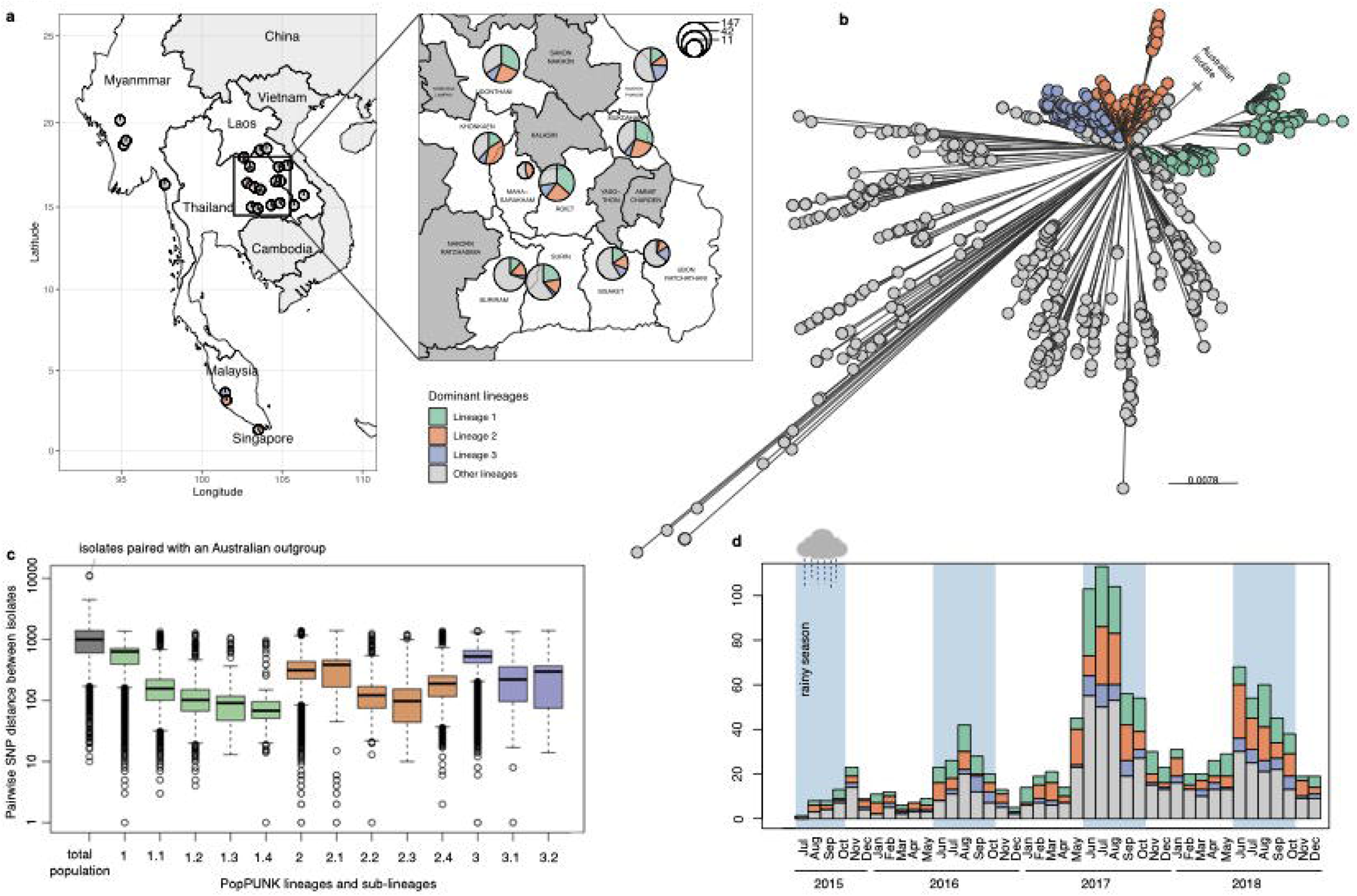
Distribution of *B. pseudomallei* genomes used in this study (a) Geographical representation of the countries and provinces sampled for the 1,391 *B. pseudomallei* genomes used in this study. Pie-chart summarises the proportion of dominant lineage 1, 2, and 3 presented at each location with the chart size proportional to the number of the samples collected (b) An unrooted phylogenetic tree colour-coded by dominant lineages (c) Histogram depicting the distribution of clinical *B. pseudomallei* isolates from the northeast Thailand cohort throughout 2015-2018 sampling period. The shaded blue area represents the period of rainy seasons. (d) Boxplots summarising the pairwised core genome SNP distances among isolates in this study, shown in a logarithmic scale. The distribution is depicted for the entire population and each dominant lineage.

## Results

### Population structure analysis revealed the successful *B. pseudomallei* lineages and mixture of clinical and environmental isolates

To define the population structure of clinical and environmental *B. pseudomallei* in a hyperendemic area of northeast Thailand and neighbouring regions (n = 1,391), we performed four independent approaches. PopPUNK^35^ analysis was performed on genome assemblies (**Supplementary data 2**). Additionally, we constructed three maximum-likelihood (ML) phylogenies^36^, each based on different sets of single nucleotide polymorphisms (SNPs): core genomes (n= 77,156 SNPs), core gene multilocus typing^37^ (cgMLST, n = 46,945 SNPs), and seven-gene multilocus typing genes^38^ (MLST, n = 31 SNPs). These approaches facilitated the grouping of isolates with close genetic similarity into distinct lineages. Notably, both PopPUNK analysis and core genome SNPs phylogeny yielded consistent results (**Supplementary Figure 1**) clustering the population into three dominant lineages (**Figure 1b**). The average pairwise core SNP distance within each dominant lineage was 549, 351, and 517 SNPs for lineage 1, 2, and 3, respectively, in contrast to the pairwise core SNP distance of 1,087 SNPs within the total population (**Figure 1c**). This lower pairwise core SNP distance across lineages confirmed the genetic relatedness as defined by PopPUNK and core genome SNP phylogenetic analysis. While cgMLST displayed conservation for two out of three dominant lineages, MLST exhibited inconsistencies across dominant lineages with lower phylogenetic resolution and poorer bootstrap support compared to other methodologies (**Supplementary Figure 1**). Consequently, we relied on the population delineated by PopPUNK and core genome SNP phylogeny for subsequent investigations.

The three predominant lineages (denoted as lineage 1 to 3) comprised 312, 297, 125 isolates, respectively. They accounted for 52.8% of the studied population and persisted throughout the sampling period. Interestingly, each lineage peaked during the rainy season, correlating agricultural practices at the onset of rainfall with increased environmental exposure and subsequent melioidosis infections (**Figure 1d**). Despite the small sample size of environmental isolates, we observed a clustering of these isolates with clinical isolates within each dominant lineage, indicating their core genetic similarities and shared origin. The ratio of environmental to clinical isolates varied across lineages (Chi-square test with Monte Carlo resampling p-value 5.00 x 10^-4^, **Supplementary Figure 2**). Due to the substantially lower number of environmental isolates used and incomplete geographical distribution matching between clinical and environmental isolates, caution is warranted in interpreting these results. Nevertheless, our findings highlight a mixing of environmental and clinical samples, suggesting that clinical isolates could serve as a surrogate for tracking the dissemination of an environmental bacterium, especially in the absence of equally comprehensive environmental samples.

### Genetic evidence identifies patterns of *B. pseudomallei* dissemination in Northeast Thailand

We examined the dissemination patterns of *B. pseudomallei* in northeast Thailand. Except for Mahasarakam where samples were limited, all three dominant lineages were present across the rest of eight studied provinces (**Figure 1a**). This prompted us to focus the analysis on these dominant lineages to identify consistent geographical distributions underlying their spread in the region. We generated lineage-specific phylogenies to improve genetic resolution for transmission analysis. Additionally, we reconstructed ancestral histories of provincial origins, quantified the number of inter-provincial transmissions to examine transmission patterns, and estimated the time of the most recent common ancestor of each dominant lineage and its sub-lineages (**Supplementary Figures 3 – 5**, see Methods). This allowed us to link the emergence of these lineages to historical events that might have affected their transmission dynamics. While transmission signals could reflect distinct dissemination patterns in each dominant lineage or its sub-lineages, we also considered the possibility of shared factors leading to a uniform geographical distribution. Notably, we observed consistent dissemination patterns in 14 out of 28 provincial pairs across the three dominant lineages (**Figure 2a, Supplementary Figure 6**). Eight out of these 14 provincial pairs potentially correlated with the slope of terrain altitude between provinces or the natural flow of rivers in the region (**Figure 2b**). These pairs included “Udon Thani-to-Khon Kaen”, “Udon Thani-to-Buriram”, “Udon Thani-to-Mukdahan”, “Udon Thani-to-Surin”, “Khon Kaen-to-Buriram”, “Nakhon Phanom-to-Mukdahan”, “Roi-Et-to-Surin” and “Surin-to-Sisaket”. Northeast Thailand is described as a saucer-shaped plateau, with elevations ranging from over 200 meters above sea level in the northwestern corner (parts of Udon-Thani, and Khon-Kaen) to less than 100 meters in the southeast (parts of Mukdahan and Sisaket), and gradually descending toward the Mekong River in the east^39,40^. The Mun River originates from elevated hills in central Thailand, streaming eastward through Buriram, Surin, Sisaket before merging with the Mekong River. Similarly, the Chi River, a tributary to the Mun, also originates from the central Thailand mountains. The Chi flows eastward through Khon Kaen, Mahasarakam, Roi-Et and converges with the Mun in Sisaket^40^.

**Figure 2.**
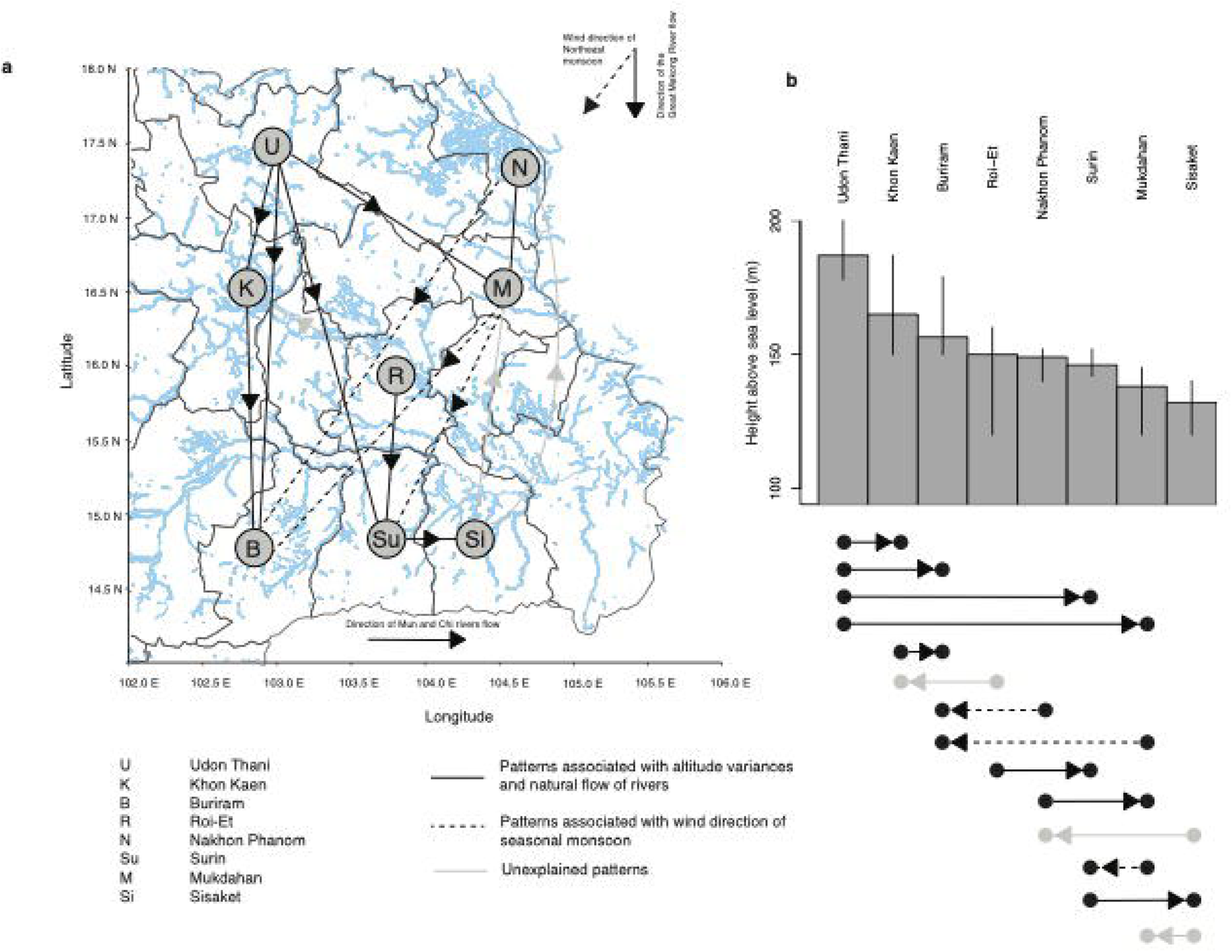
Dissemination patterns in northeast Thailand. (a) Province-to-province transmission patterns influenced by northeast Thailand geographical landscape. Nodes present provinces, denoted by abbreviation and ordered by altitude: U-Udon Thani, K-Khon Kaen, B-Buriram, R – Roi Et, N-Nakhon Phanom, Su-Surin, M-Mukdahan, and Si-Sisaket. Rivers are depicted in blue with major rivers including the Great Mekong River, the Chi River, and the Mun River and their flow direction annotated. (b) Average altitude of provinces in meters above sea level. Error bars present 95% confidence interval. The northwest provinces exhibit higher altitudes, gradually declining towards the southeast. For (a) and (b) solid arrows illustrating transmission directionality explained by altitude differences. Dotted arrows represent transmission directionality influenced by northeast monsoon winds. Grey arrows signify patterns with unclear explanation.

Additionally, another set of three out of 14 conserved patterns coincided with the wind direction of the northeastern monsoon during the dry season (“Nakhon Phanom-to-Buriram”, “Mukdahan-to-Buriram”, and “Mukdahan-to-Surin”). Thailand experiences two predominant monsoon seasons: the southwest monsoon from May to October and the northeast monsoon from November to April. The southwest monsoon brings heavy rainfall from the southwest to northeast, often marking the start of the agriculture season (**Figure 1c**)^41,42^. Conversely, the northeast monsoon brings dry winds from the northeast to southwest. Our observation implies the potential for dry winds to transport aerosolised soil contaminated with *B. pseudomallei* westward. Despite that previous air sampling during Thailand’s rainy season did not detect *B. pseudomallei*, exploring the impact of dry winds during the northeastern monsoon is essential. This consideration becomes even more pertinent given reports of the long-range transport of particles and small organic matters, such as PM2.5 and PM10, via the northeast monsoon elsewhere in Southeast Asia^43,44^. Furthermore, we successfully estimated the time of most recent of ancestry for a sub-lineage 1.3, a descendant of lineage 1. Our finding revealed that this sub-lineage emerged around 2011 (95% HPD of 2000-2014, **Supplementary Figure 3**). The age of this sub-lineage implies that older lineages, such as its parent lineage 1 likely experienced multiple monsoon seasons, which possibly resulted in the observed patterns. Apart from the terrain slope, inland rivers, canal systems^45^ and regular monsoons^41,42^; various factors likely contributed to shaping *B. pseudomallei* dissemination in northeast Thailand. Anthropogenic activities, such as human migration between the provinces, may also contribute to the observed pattern; however without access to comprehensive human movement data, this aspect remains challenging to investigate.

### Genetic markers potentially contributing to the emergence of successful *B. pseudomallei* lineages

The co-existence of multiple *B. pseudomallei* lineages within the same geographical areas and timeframe implies the presence of diverse adaptive strategies which enable them to thrive in a shared ecological niche. While some smaller lineages may be sporadically detected, the persistence of the three dominant lineages throughout the sampling period supports their fitness and successful adaptive strategies in this niche. We next sought to identify genes that were present in isolates that form each dominant lineage, or its sub-lineage; but absent in non-dominant lineages (see Methods). Out of total 15,237 genes in the pan-genome outlined from this population (see Methods), 5,577 genes were conserved across the entire population while 9,660 genes were variably present (accessory genes). Dominant lineage-specific genes were defined as accessory genes present in ≥95% of isolates within any of the dominant lineages or their sub-lineages, but present in ≤15% of isolates outside these lineages. Among these, 247 genes were identified as lineage-specific with their specificity to each dominant lineage and sub-lineage tabulated in **Supplementary data 3**. The majority of dominant lineage-specific genes were poorly characterised and annotated as hypothetical proteins (**Figure 3**). To gain insights into the potential functions, we annotated them using Gene Ontology (GO terms)^46^ which classify them by Biological process, Molecular function, and Cellular component (Figure 3b, see Methods). Of the 247 dominant lineage genes, GO terms could be assigned to 27 genes for Biological Process, 68 for Molecular Function, and 12 for Cellular component. For genes that could be assigned GO terms, functions involved in “DNA integration”, “DNA recombination”, and “DNA methylation” might indicate their potential roles in horizontal gene acquisition and protection against incoming foreign DNA through site-specific DNA methylation. Furthermore, GO terms associated with “DNA binding” and “Regulation of DNA-templated transcription” may suggest lineage-specific regulation of the expression of these genes.

**Figure 3.**
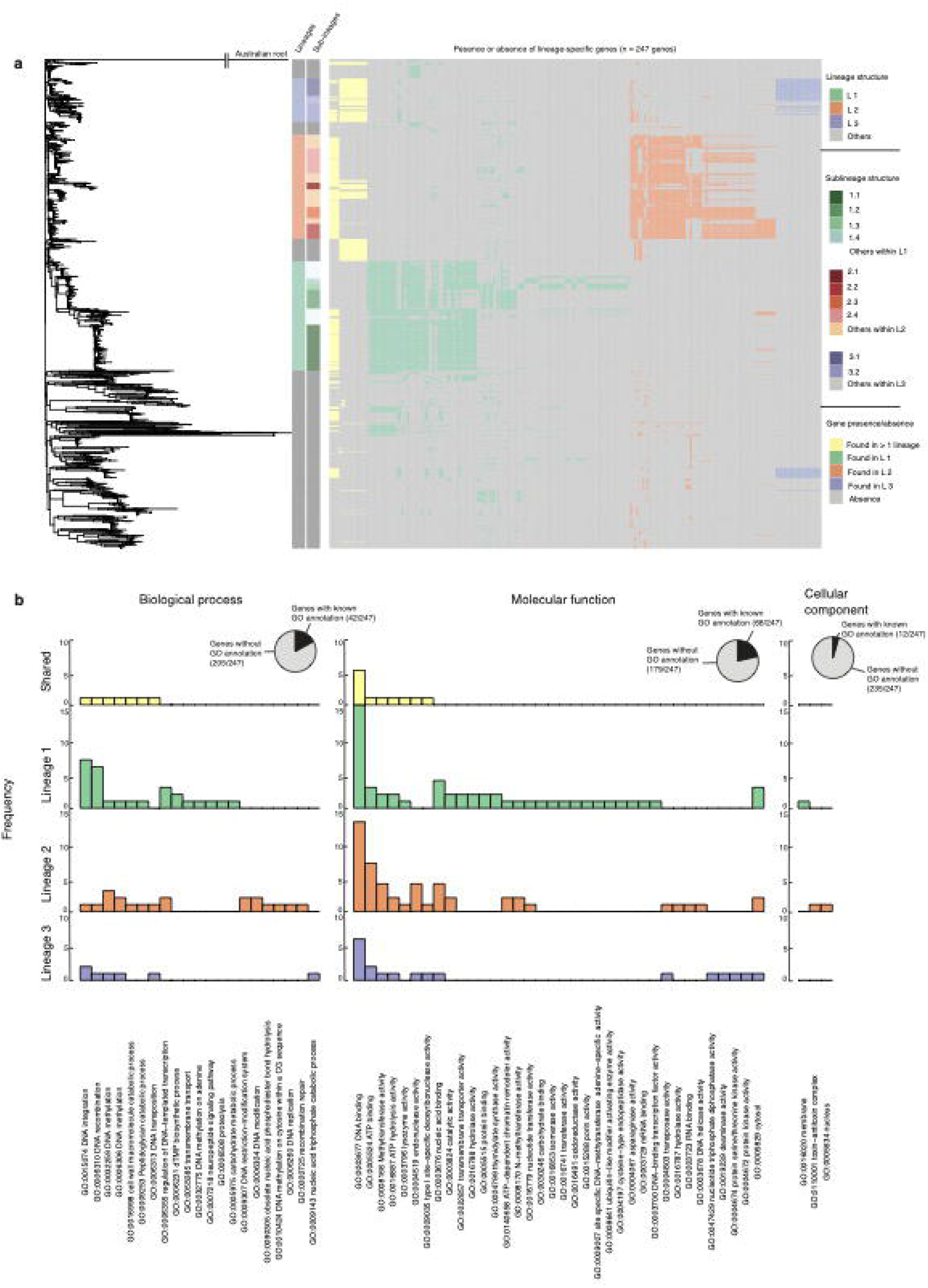
Dominant lineage-specific genes and their Gene Ontology (GO terms). (a) The heatmap represents lineage-specific genes (right) detected in each isolate, aligned with the phylogeny (left). Lineage-specific genes shared across multiple dominant lineages are highlighted in yellow. Lineage-specific genes from lineage 1, 2, 3 are coloured in green, red, and purple, respectively. Additionally, the colour stripes provide information on the lineage and sub-lineage membership (b) Bar plots displays the frequency of GO annotations of lineage-specific genes in each dominant lineage categorised by biological process, molecular function, and cellular compartment. The pie-charts summarise the proportion of lineage-specific genes with assigned GO terms (black).

### Lineage-specific genes were selectively expressed

To delve deeper into the functionality of dominant lineage-specific genes, we explored their expression patterns across both environmental and infection conditions. We selected representative strains – “K96243” (lineage 1 – sub-lineage 1.1), “UKMD286” (lineage 2 – sub-lineage 2.1), and “UKMH10” (lineage 3 – sub-lineage 3.2) – all are clinical isolates with pre-existing gene expression profiles under infection and environmental conditions^47–49^. For environmental conditions, K96243 was exposed to water^47^, while UKMD286 and UKMH10 were cultivated in a soil extract medium^48,49^ to mimic *B. pseudomallei* in the environment. For infection conditions, K96243 and UKMD286 were used in murine challenges^47,48^ and isolated from mice organs, while UKMH10 was subjected to human plasma^49^ to simulate host infection. This approach facilitated the comparison of differentially expressed lineage-specific genes between environmental and infection conditions. Although each representative strain carried a complete set of lineage-specific genes for their respective sub-lineages (K96243 with 47 genes, UKMD286 with 27 genes, and UKMH10 with 14 genes), their collective representation accounted for 69 out of 247 total lineage-specific genes (27.9%) due to observed genetic diversity within the dominant lineage. Notably, 11 out of 47 lineage-specific genes in K96243 and 6 out of 27 lineage-specific genes in UKMD286 were up-regulated in the environmental conditions (**Figure 4a to 4c, Supplementary data 4**). Notably, none of the lineage-specific genes showed up-regulation during the infection condition. The remaining lineage-specific genes did not exhibit preferential expression in either environmental or infection conditions. The elevated expression level of lineage-specific genes in the environmental condition was unexpected considering that all representative strains were clinical isolates. This observation potentially suggests that dominant lineage-specific genes may play more substantial roles in bacterial environmental survival than in host pathogenicity.

**Figure 4.**
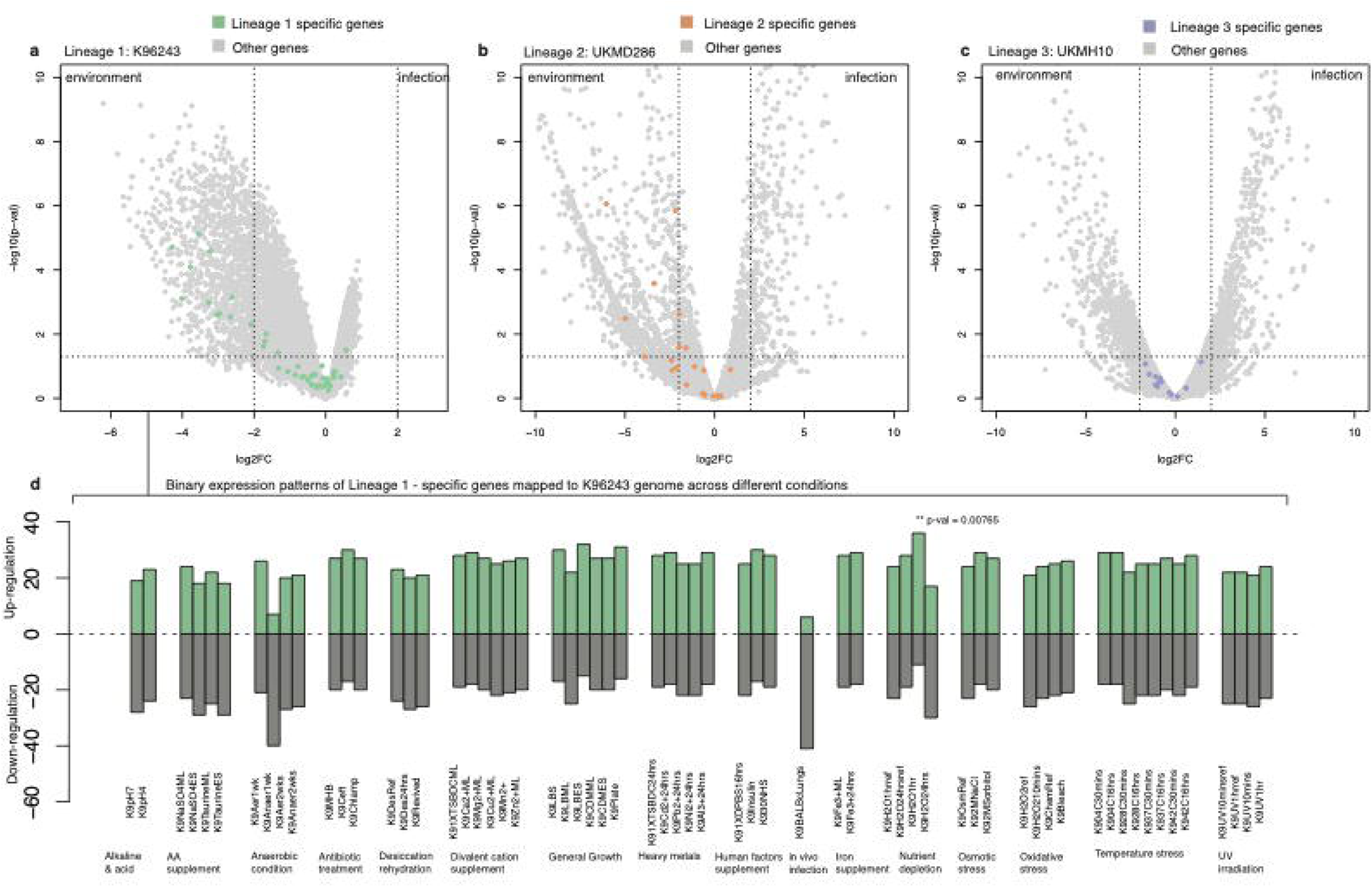
Transcriptome analysis of representative strains: K96243 (lineage 1), UKMD286 (lineage 2) and UKMH10 (lineage 3) (a to c) Volcano plots demonstrate differential gene expression (DGE) between environmental and infection conditions. Vertical dotted lines represent the statistical cut-off at log two-fold change, while horizontal dotted lines display the statistical cut-off at the adjusted p-value of 0.05 on a negative log scale. Each dot represents a gene, with lineage-1, lineage-2, and lineage-3-specific coloured in green, red, and purple, respectively. (d) Binary expression profile of lineage-1-specific genes across different conditions. A star denotes significant differences in the gene expression profile of lineage-1-specific genes compared the remaining genes of strain K96243.

To thrive in its environmental habitat, *B. pseudomallei* must cope with ranges of physical, chemical and biological stresses such as desiccation, temperature fluctuations, osmotic changes, oxidative stress, UV exposure, nutrient scarcity, changes in pH, exposure to heavy metals, competition from antibiotics released by other microbes, and predation by eukaryotes. We leveraged the extensive condition-wide transcriptome data spanning 62 conditions available for K96243^47^, a representative strain from lineage 1, to compare the expression patterns of lineage-1-specific genes with the rest of genes in the K96243 genome (**Figure 4d**). Our analysis revealed that lineage-1-specific genes within K96243 exhibited a higher level of gene expression when the bacterial cell experienced nutrient deprivation compared to other genes in K96243 genome (Two-sided Fisher’s exact test p-value = 9.23 x 10^-5^). This finding implies that lineage 1 might possess an adaptive strategy to persist in nutrient-depleted soil, which is not uncommon in melioidosis endemic areas^27,28,50^, before being acquired by a human host and subsequently causing the disease.

### Example of lineage-specific genes

The majority of lineage-specific genes were located within genomic islands (GI)^30,51,52^. These regions are characterised by anomalies in %G+C content or dinucleotide frequency signatures, or the presence of genes associated with mobile genetic elements such as insertion sequence (IS) elements and bacteriophages. Notably, we observed that a cluster of genes specific to lineage 1 (*BPSS2060* to *BPSS2072*) formed a mosaic structure within a putative metabolic island known as GI 16^30^. Although several variations of GI 16 have been reported (GI16, GI16.1, GI16.2, GI16a, GI16b, and GI16b.1)^51^, it typically spans 60 kb (*BPSS2051* to *BPSS2090*) and carries several known virulence determinants and genes that enhance metabolic versatility. While certain virulence factors, such as the filamentous haemagglutinin (*BPSS2053*) required for host cell adhesion and its processing protein were conserved across multiple lineages observed in our study, genes encoding functions that potentially expand the metabolic repertoire were specific to dominant lineages. For example, the mosaic structure of GI 16 (*BPSS2060* to *BPSS2072*), specific to lineage 1 (**Supplementary data 3**), contains genes involved in alternative nutrient catabolism and anabolism (*BPSS2060*, *BPSS2065*, *BPSS2067*, *BPSS2068*, and *BPSS2072*), transcriptional regulation (B*PSS2061*), and substrate transport (*BPSS2064*, *BPSS2071*). Out of 11 lineage-specific genes located in the mosaic structure of GI 16, eight were found to be upregulated during the early phase of nutrient starvation while remaining silent during infections. This finding reflects the functional division of GI16 where its lineage-specific mosaic structure contains genes that contribute to metabolic versatility, while its core structure encodes virulent determinants associated with disease implications. It is important to note that the structure of GI 16 may vary across different regions due to the plasticity of genomic islands and changes in selection pressures. While this observation is significant for a dominant lineage in northeast Thailand, it may not be generalisable to other geographical locations.

### Lineage-specific genes were introduced by homologous recombination

Homologous recombination has been shown to play a significant role in facilitating the acquisition and loss of genes, and the generation of mosaic structures within the GI of *B. pseudomallei*^53^. To better understand its association with lineage-specific genes, we identified recombination events and quantified the rates of recombination in the dominant lineages. The ratio of polymorphisms introduced through recombination compared to those introduced by mutation (*r/m*) was 3.7, 4.6 and 2.2 for lineages 1, 2, and 3 respectively (**Table 1**). A very high proportion of genes underwent recombination at least once: 99.5% of genes in lineage 1, 99.9% in lineage 2, and 96.6% in lineage 3. Furthermore, every lineage-specific gene within each dominant lineage underwent recombination (**Supplementary Figure 8**). The bacterial restriction modification (RM) systems prevent the invasion of foreign DNA and restrict gene flow between *B. pseudomallei* lineages^31^. Notably, components of this system including a type I restriction system and modification methylase were among dominant lineage-specific genes (*BPSL0947-BPSL0948* in lineage 1, and their homologues in lineage 2 and 3). They may act as a barrier for homologous recombination and potentially modulate lineage-specific genetic diversity. This highlights the intricate interplay between recombination, lineage-specific genes, and the RM system in shaping the genetic landscape of *B. pseudomallei* in northeast Thailand.

**Table 1.**
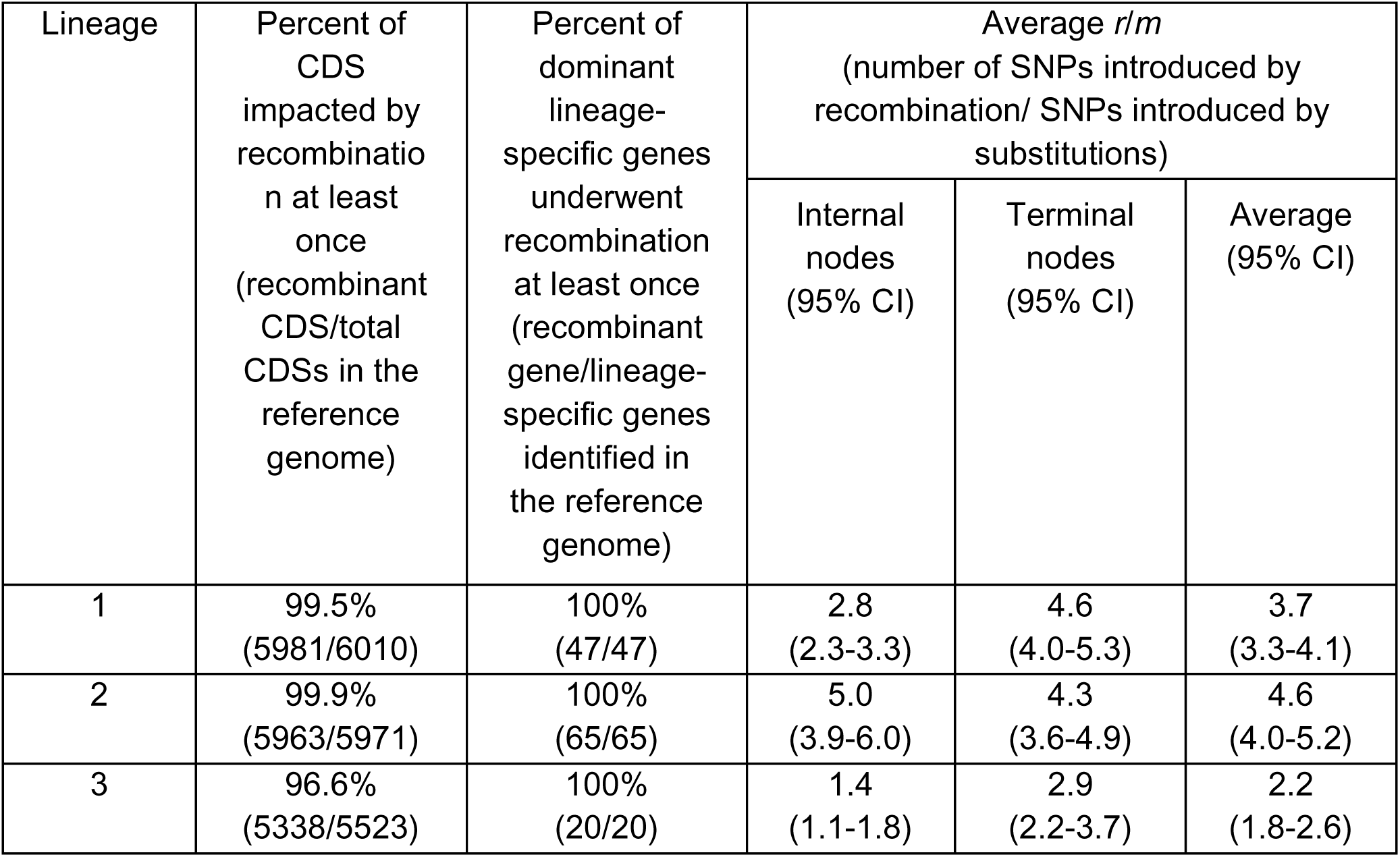
Recombination in dominant lineages.

| Lineage | Percent of CDS impacted by recombination at least once (recombinant CDS/total CDSs in the reference genome) | Percent of dominant lineage-specific genes underwent recombination at least once (recombinant gene/lineage-specific genes identified in the reference genome) | Average *r/m*<br>(number of SNPs introduced by recombination/ SNPs introduced by substitutions) |  |  |
| --- | --- | --- | --- | --- | --- |
|  |  |  | Internal nodes<br>(95% CI) | Terminal nodes<br>(95% CI) | Average<br>(95% CI) |
| 1 | 99.5%<br>(5981/6010) | 100%<br>(47/47) | 2.8<br>(2.3-3.3) | 4.6<br>(4.0-5.3) | 3.7<br>(3.3-4.1) |
| 2 | 99.9%<br>(5963/5971) | 100%<br>(65/65) | 5.0<br>(3.9-6.0) | 4.3<br>(3.6-4.9) | 4.6<br>(4.0-5.2) |
| 3 | 96.6%<br>(5338/5523) | 100%<br>(20/20) | 1.4<br>(1.1-1.8) | 2.9<br>(2.2-3.7) | 2.2<br>(1.8-2.6) |

## Discussion

Our analysis of *B. pseudomallei* population genomics enhances our understanding of the evolution and adaptive strategies employed by the dominant lineages in the melioidosis hyperendemic region of northeast Thailand and neighbouring countries. Through an unprecedentedly dense sampling effort between 2015 and 2018, we were able to determine the co-existence of three dominant lineages, characterise their dissemination patterns, and identify lineage-specific genes possibly contributing to their success during the studied period. By analysing transcriptome data from representative strains of each dominant lineage, we gained further insights into their adaptive strategies, particularly emphasising bacterial persistence in the environment as crucial for subsequent host acquisition and infection. Nevertheless, our study has a few limitations.

Due to limited environmental surveillance and the scarcity of environmental isolates in Southeast Asia, our study primarily relied on clinical isolates. Nonetheless, the co-occurrence of clinical and environmental isolates within the same lineage, coupled with the direct acquisition of clinical isolates from the environment, suggests that our findings may hold broader implications for environmental isolates. Furthermore, the lineage classification report in our study is subject to potential alterations over time with the introduction of new data. As *B. pseudomallei* continuously adapts to environmental pressures, the composition of lineages 1, 2, and 3, along with their respective lineage-specific genes may shift. New lineages with superior fitness or selective advantages, of different adaptive strategies could potentially outcompete the existing dominant lineages. Consequently, this may lead to emergence of new lineage classifications in the future. Continued surveillance efforts including both clinical and environmental samples will be essential. This ongoing monitoring will be pivotal in identifying alterations within the bacterial population, uncovering new adaptive strategies, and evaluating their impact on disease dynamics over time.

Our understanding of the significance of bacterial persistence as a strategy for successful lineages stems from the observed up-regulation of dominant lineage-specific genes in environmental conditions, along with the increased expression of lineage-1-specific genes during nutrient deprivation. However, it is essential to recognise the genetic diversity within each dominant lineage. The representative strains used in our study carried a subset of the lineage-specific genes corresponding to their lineage. As a result, lineage-specific genes absent in these strains, often annotated as hypothetical proteins, remained unexplored in our analysis. Moreover, our detailed characterisation of lineage-1-specific genes using a comprehensive transcriptome dataset of 62 distinct conditions might not capture the full spectrum of conditions encountered by *B. pseudomallei* in its natural habitat. There is a possibility that other adaptive strategies were overlooked in this study, indicating scope for future exploration. Despite these shortcomings, our dataset and analysis currently represent one of the most comprehensive efforts to date. Future research will prioritise the generation of a more extensive condition-wide transcriptome, covering a broader range of conditions and incorporating strains with diverse genetic variations to identify other adaptive strategies employed by *B. pseudomallei*.

Our findings underscore the significance of environmental persistence in driving the success of dominant lineages 1 and 2, notably highlighting lineage-1-specific genes in mediating bacterial survival under nutrient depletion. It remains uncertain whether lineage 3 adopts a similar strategy. Nevertheless, our results align with previous soil sampling studies, which consistently observed a higher prevalence of *B. pseudomallei* in nutrient-depleted compared to nutrient-rich soil^28,50^. Additionally, a molecular evolutionary study also supports the species’ long-term adaptation to survive nutrient scarcity^27^. The northeast region of Thailand, where our samples were primarily collected, has inherently low-fertility soil. This is exacerbated by intensified agriculture, monoculture, excessive synthetic fertilizer use, and poor land management, resulting in depleted soil nutrients and organic matter^54^. This presents a challenging environment for *B. pseudomallei* to thrive, thereby potentially selecting for successful lineages with persistent traits as observed in our study.

Our analyses also highlight the role of various factors such as differences in terrain altitude^55,56^, river flow dynamics^40^, and the northeast monsoon^41^ contribute to shaping the dissemination patterns of *B. pseudomallei*. These drivers of dissemination are influenced by both natural and human activities. For instance, strong winds can carry dried soil particles^57^, potentially containing *B. pseudomallei* over distances. Climate change-induced alterations in vegetation cover might expose soil to rainfall and winds^58^, impacting the bacterial spread. Additionally, deforestation can disrupt natural barriers like trees and shrubs, accelerating water runoff^57,58^ and potentially facilitating the wide-ranging dissemination of *B. pseudomallei* during flood. Considering these dynamics, the strategy of bacterial persistence likely plays a pivotal role in its widespread dissemination within the region, thereby influencing disease prevalence. Therefore, an effective disease control strategy should integrate both environmental and clinical public health measures to effectively mitigate the impact of melioidosis.

## Materials and Methods

### Data collection and bacterial isolates

The *B. pseudomallei* isolates in our study included a cohort^8^ from northeast Thailand gathered between July 2015 to December 2018 consisting of 1,265 clinical isolates. We also incorporated a contemporaneous dataset from Thailand and neighbouring regions^15,29–34^, comprising 15 clinical and 111 environmental isolates sourced from previous publications. In total, 1,391 *B. pseudomlalei* genomes were used in this study. Their metadata and accession numbers were documented in **Supplementary data 1**.

The northeast Thailand collection was collected from patients participated in our longitudinal cohort study^7^. The patients were from nine provinces including Udon Thani (n = 230), Mukdahan (n = 198), Roi Et (n = 195), Surin (n = 170), Nakhon Phanom (n = 135), Buriram (n = 123), Sisaket (n = 107), Khon Kaen (n = 96) and Maha Sarakham (n = 11), who were admitted to nine hospitals included in our cohort. The ethical approval for the cohort study was obtained from the Ethics Committee of the Faculty of Tropical Medicine, Mahidol University (MUTM 2015-002-001 and MUTM 2021-055-01). The isolates were obtained from various clinical samples, including blood (71.8%), pus (12.5%), sputum (9.9%), body fluid (3.5%), urine (1.9%) and tissue (0.4%) (**Supplementary data 1**). As latent infection accounts for < 5% of the cases, the majority of clinical cases likely directly acquired from the environment^8^. The numbers of enrolled cases were lower at the beginning of the study due to the delayed sample collection in some study sites, resulting in inconsistent number of bacterial isolates used across different sites. When applicable, a permutation test was performed to ensure that an unequal number of isolates did not impact the temporal or spatial analysis.

### Culture confirmation of *B. pseudomallei*, DNA extraction and whole genome sequencing

All 1,265 *B. pseudomallei* samples from the northeast Thailand collection were cultured on Ashdown, selective agar plates and confirmed the species using latex agglutination test and matrix-laser absorption ionisation mass spectrometry (MALDI-TOF MS). A single colony from Ashdown agar plate was subjected to culture in Luria-Bertani (LB) broth and subsequently used for DNA extraction. Genomic DNA was extracted using QIAamp DNA Mini Kit (Qiagen, Germany). All genomic DNA were processed for the 150-base-read library preparation and sequencing using Illumina HiSeq2000 system with 100-cycle paired-end runs at Wellcome Sanger Institute, Cambridge UK. An average of 71X read depth was achieved. To control the potential contamination in each sample with other closely related species, we assigned taxonomic identity using Kraken^59^ v.1.1.1. We then estimated the genome completeness and species confirmation using CheckM^60^ v.1.2.2 and FastANI^61^ v.1.31, respectively. The quality control data of the 1,265 genomes were listed in **Supplementary data 2**.

### Genome assembly and mapping from short read data

Short reads were *de novo* assembled using Velvet v.1.2.10^62^ followed by optimisation as previously described^29^, with their quality scores reported in **Supplementary data 2**. Short reads were also mapped against several reference genomes, including a strain K96243^30^ (accession numbers BX571965 and BX571966) to determine the whole population structure, and lineage-specific references to improve the resolution for lineage-specific analyses. We selected K96243^30^ genome as a population-wide reference due to its origin in northeast Thailand, aligning with the geographical focus of our study. Additionally, its well-characterised and complete genome further streamlined subsequent analyses. For all mapping, variants were called using Snippy v.4.6.0 (https://github.com/tseemann/snippy). To avoid mapping errors and false SNPs, we filtered out SNPs covered by less than 10 reads and found in a frequency of less than 0.9.

### Defining whole population structure

#### PopPUNK clustering

PopPUNK v.2.6.0^35^ was run on 1,391 assembled genomes. To define the core and accessory distance between each pair of isolates, the assemblies were hashed at different k-mers. The population model was fit using command line “poppunk-runner.py --fit-model –distance <database.dists> --output <database> --full-db --ref-db <database> --min-kmer 15 –max-kmer 31 --max-a-dist 0.53 --K 4 –k-step 2” with the result density of 0.028, transitivity of 0.992, and network score of 0.8961.

#### Maximum likelihood phylogenies from core genome SNP, cgMLST and MLST

An alignment of full genome was created by mapping whole genome sequences of each *B. pseudomallei* against a complete genome of K96243^30^ strain. From this alignment, 4,221 cgMLST loci based on a scheme described in^37^ were extracted and concatenated to form cgMLST alignment. Additionally, seven MLST loci, as per scheme described in ^38^ were extracted from the same alignment and concatenated to create the MLST alignment. Core genome SNP alignment was identified from a full genome alignment using snp-sites^63^ v.2.5.1, with genomic islands^51^ masked. Separate maximum likelihood phylogenies were constructed for core genome SNP alignment, cgMLST alignment, and MLST alignment using IQ-TREE^36^ v.2.0.3. Standard model selection in IQ-TREE determined the best-fit model as TVM+F+ASC+R6 for all three phylogenies. To access the robustness of the phylogenetic trees, a 1,000 bootstrap support was performed for each tree.

#### Comparison of population structure outlined phylogeny constructed from core genome SNPs, cgMLST, MLST and PopPUNK

To test for consistency between phylogenetic trees constructed from core genome SNPs, cgMLST and MLST alignment, we use the R package treespace^64^ v. 1.1.4.3 to explore the tree tip distributions. We compared pairwise tree distances within the first 100 bootstraps within each alignment category (indicative of bootstrap support strength), and across trees generated from different alignment categories (indicating proximity between tree categories). The tree pairwise distances were computed, and principal components (PCs) were derived with eigenvalues calculated for different PCs. The similarity among phylogenies from each alignment category was assessed using two PCs dimensions, which jointly accounted for >90% of variability in pairwise distance (**Supplementary Figure 1a**). The scatter plot of PCs revealed a close clustering of bootstrap trees from core genome SNP and cgMLST, while the bootstrap trees from MLST alignment showed greater dispersion, highlighting less consistency in the trees generated by the MLST approach. We further compared the consistency between the median phylogenetic tree of each alignment category and PopPUNK classification was visually compared using iTOL^65^ (**Supplementary Figure 1b**).

#### Specific lineage analysis

To investigate the dissemination and genetic diversity within each dominant lineage, we conducted individual genome alignments, recombination removal, and maximum likelihood phylogeny. To enhance the sensitivity of variant, we selected closely related genomes as references for each lineage. Specifically, the complete genome *B. pseudomallei* strain K96243 (accession numbers BX571965 and BX571966) served as the mapping reference for lineage 1, while new reference genomes were created for lineages 2 and 3.

For lineage 2, we chose a representative isolate 27035_8#57 and subjected it to long-read sequencing on a local MinION sequencer following the manufacturer’s standard protocol (Oxford Nanopore Technologies, Oxford, United Kingdom). A complete hybrid assembly of the long-read and short-read sequence data of this strain was performed using Unicycler^66^ v.0.8.4.

The resulting hybrid assembly of the 27035_8#57 genome was employed as the mapping reference for this lineage.

In the absence of representative complete genomes for lineage 3, we selected the best quality de novo assembly of isolate 27035_8#119 and orientated its contigs according to strain K96243 using ABACAS^67^ v.1.3.1. This genome was used as a mapping reference for lineage 3.

For lineage-specific mapping, Snippy v.4.6.0 as employed as in the whole population analysis. All genome alignments were subjected to Gubbins^68^ v.3.1.3, a recombination identification tool, to detect and remove recombination fragments. This process determined the genetic diversity introduced by horizontally acquired elements and vertically inherited SNPs, thereby producing recombination-free SNP alignments for phylogenetic reconstruction. Maximum-likelihood phylogenies were constructed using recombination-free SNP alignment of each dominant lineage using IQ-TREE^36^ v.2.0.3 with TVM+F+ASC+R6 and 1,000 replicates of bootstrap support. The overall proportion of nodes with ≥80% bootstrap support of lineage-specific phylogenies reached 83.5%.

#### Dating the timeline for lineages and sub-lineages

To enable dating analysis, we further divided each lineage into sub-lineages using R package rhierbaps^69^ v.1.1.4. We inferred evolutionary timeline and estimated the age of each lineage and sub-lineage based on the isolate’s collection date. A Bayesian molecular dating provided in R package BactDating^70^ v.1.1.1. was employed to assess the temporal signals by examining a positive correlation between the isolate’s collection date and the root-to-tip distance. Recombination removed phylogenies were used in this analysis. A date-randomisation test, consisting of 100 permutations, was performed to assess the robustness of the temporal signal compared to noise.

Notably, the temporal signals were discernable at the sub-lineage level rather than the broader lineage level. Among the 10 sub-lineages, only one sub-lineage (lineage 1.3) exhibited a positive correlation in their clock signals. Given the limited sample size of lineage 1.3, we employed a strict clock model to prevent parameter over-fitting. We ran three independent Markov chain Monte Carlo (MCMC) chains, each spanning at least 100 million iterations, and sampled every 10,000 steps. The prior mutation rate derived from Pearson and colleagues was used. Visual inspection of the trace from each MCMC chain confirmed signal convergence, with effective sampling size values > 200 for key parameters. Visualisation of results were performed using the R package ggtree^71^ v.3.10.0 to generate credibility time-calibrated phylogeny for each sub-lineage (**Supplementary Figures 3**).

#### Ancestral state reconstruction analysis

Ancestral trait reconstruction was conducted to discern the dissemination patterns of *B. pseudomallei* among provinces in northeast Thailand, focusing on dominant lineages. Due to varying number of isolates among provinces, the analysis excluded Mahasarakham, which had a limited dataset (n = 11), resulting in the analysis of eight provinces: Buriram, Khon Kaen, Mukdahan, Nakhon Phanom, Roi Et, Sisaket, Surin, and Udon Thani. This approach yielded 28 potential province-to-province transmission combinations. To mitigate sampling biases, we sub-sampled the phylogeny of each dominant lineage to have an equal number of isolates per province (n = 15 isolates) and permuted 1,000 times. Using the stochastic character mapping function (*make.simmap*) from the R package phytools^72^ v.1.9.16, we conducted 100 simulations (nsim = 100) to reconstruct the provincial origins at each node in the sub-sampled phylogeny (1,000 phylogenies per lineage). This allowed us to quantify transition events (Markov jumps) between province pairs and determine the cumulative branch length associated with each province (Markov rewards). A Mann-Whitney U test, with Bonferroni correction for multiple comparisons was applied to compare the transition event counts among provinces (**Supplementary Figure 6**).

#### Pan-genome analysis

All the study genomes were annotated using Prokka^73^ v.1.14.5, and further used in the pan-genome analysis. Each genome has a median of 5,845 coding sequences (CDS) predicted onto each genome with a range of 5,642 to 6,142 CDS per genome. Panaroo^74^ v.1.3.3 was employed to estimate the pan-genome with a sensitive option and a cut-off sequence identity of 92% derived from previous study^15^. The number of estimated genes falls within a comparable range to previous studies from a single population^29,53^.

#### Identification of dominant lineage-specific genes

We determined lineage-specific genes by assessing their prevalence within dominant lineage or any of their sub-lineages, requiring a high occurrence (95%) within these specific groups while maintaining a low presence in non-dominant lineages. To achieve this, three thresholds were employed: strict (95 % occurrence in dominants vs 5% occurrence in non-dominants), intermediate (95% in dominants vs 10% in non-dominants), and relaxed (95% in dominants vs 15% in non-dominants). Based on visual examination of gene distribution patterns (**Figure 3a, Supplementary** Figure 7), the relaxed threshold was used to maximise the number of genes included in subsequent analysis.

#### Identification of Gene Ontology (GO: terms)

Amino acid sequences of lineage-specific genes were submitted to InterPro database^46^ (https://www.ebi.ac.uk/interpro/) which characterised the function of lineage-specific genes based on biological processes, molecular functions, and cellular compartments **(Figure 3; Supplementary data 3**).

### Transcriptomic analysis of dominant lineage-specific genes

#### Lineage 1 transcriptome analysis

The analysis focused on an expression profile of a strain K96243, which serves as a representative of lineage 1. Data was sourced from microarray experiment generated by Ooi *et al*.^47^ and accessed through the Gene Expression Omnibus (GEO) under accession number GSE43205.

To understand the difference between environmental and infection conditions, we compared the expression profile of K96243 being exposed to water and K96243 recovered from infected mice. To simulate environmental conditions, K96243 was cultured to log phase in LB medium, subsequently washed with sterile deionising water, and suspended in water. To emulate infection conditions, BALB mice were infected with 1000 CFU of *B. pseudomallei*, and bacteria were harvested from the lungs three days post-infection. Two replicates were performed for each condition. We retrieved microarray data from GEO using the R package GEOquery^75^ v.2.58.0, and differential gene expression analysis was performed using the R package limma^76^ v.3.58.1.

We used binary expression patterns reported in Ooi *et al*.^47^ to compare the expression profiles of lineage-specific genes against the remaining genes in K96243 across 62 conditions. This enabled the comparison of the count of expressed genes within the lineage-specific category against the remainder of the genes for each condition using Fisher’s exact test, with multiple testing adjustments via Benjamini-Hochberg corrections.

#### Lineage 2 transcriptome analysis

This analysis focused on the expression profile of a strain UKMD286, representative of lineage 2. RNAseq data was obtained from an experiment conducted by Ghazali *et al.* ^48^ and accessed through the European Nucleotide Archive (ENA) (E-MTAB-11200).

To simulate environmental conditions, UKMD286 was cultured in BHIB medium overnight, resuspended, and inoculated into soil extract medium. For infection condition, BALB mice were infected with UKMD286, and the bacteria were harvested from spleens five days post infection. Each experiment was conducted with two replicates. FastQC v.0.11.9 and FastXtool v.0.0.14 were used to pre-processed sequenced reads. Raw reads were aligned to UKDM286 genome using Hisat2^77^ v. 2.2.1 with differential gene expression performed using the R package DESeq2^78^ v.1.40.2.

#### Lineage 3 transcriptome analysis

We used a strain UKMH10 to represent the expression profile of lineage 3. Data was originated from an RNAseq experiment conducted by Kong *et al.*^49^ and was accessed through the European Nucleotide Archive (ENA) (PRJEB53338)

To replicate environmental conditions, UKMH10 was cultured in LB medium overnight and sub-cultured into soil extract medium. To simulate infection conditions, UKMH10 was cultured in LB medium overnight and inoculated into human plasma, then incubated at 37 C to mimic the human body temperature. Bacterial cells were harvested once the absorbance reading of the bacterial cultures at 600 nm (OD600) reached 0.5. Each experiment was performed with two replicates. FastQC v.0.11.9 and FastXtool v.0.0.14 were used to pre-processed sequenced reads. Raw reads were aligned to UKMH10 genome using Hisat2^77^ v.2.2.1 with differential gene expression performed using the R package DESeq2^78^ v.1.40.2.

## Acknowledgements

We would like to thank physician and microbiology staff at the Udon Thani Hospital, Khon Kaen Hospital, Srinakarin Hospital, Nakhon Phanom Hospital, Mukdahan Hospital, Roi Et Hospital, Surin Hospital, Sisaket Hospital and Buriram Hospital for reporting the case and providing the facilities for bacterial collection. We sincerely appreciate the transcriptomic data of B*. pseudomallei* UKMD286 and UKMH10 provided by Professor Sheila Nathan and colleagues. We are grateful to Dr. Soe Htet Aung for supporting geographical map plot. This work was funded by Mahidol University (MU-KMUTT Biomedical engineering and Biomaterials Consortium) and Royal Golden Jubilee Ph.D. Programme (RGJ-ASEAN) (http://rgj.trf.or.th) through RS and NC. CCho was funded by Wellcome International Master Fellowship (221418/Z/20/Z). CChe was funded by Wellcome International Intermediate Fellowship (216457/Z/19/Z) and Sanger International Fellowship. NC and TEW were supported by the National Institute of Allergy and Infectious Diseases of the National Institutes of Health (NIAID/NIH) (https://www.nih.gov) under Award Number U01AI115520. The content is solely the responsibility of the authors and does not necessarily represent the official views of the National Institutes of Health. This research was funded in part, by the Wellcome Trust [220211]. For the purpose of Open Access, the author has applied a CC BY public copyright license to any Author Accepted Manuscript version arising from this submission. The funders had no role in study design, data collection and analysis, decision to publish, or preparation of the manuscript.

## Data availability

The genome sequence data presented in this study can be found in online repositories. The ENA under study accession number PRJEB25606 and PRJEB35787. The accession numbers for individual genomes, and their annotated assemblies are listed in **Supplementary data 1**.

## Code availability

The analyses used public software including Kraken v.1.1.1, CheckM v.1.2.2, FastANI v.1.31, FastQC v.0.11.9, FastXtool v.0.0.14, Unicycler v.0.8.4, ABACAS v.1.3.1, Velvet v.1.2.10, Prokka v.1.13.4, Panaroo v.1.2.9, Snippy v.4.6.0, PopPUNK v.2.6.0, IQ-TREE v.2.0.3, Gubbins v.3.1.3, Hisat2 v.2.2.1 and R packages: treespace v.1.1.4.3, BactDating v.1.1.1, ggtree v.3.10.0, phytools v.1.9.16, GEOquery v.2.58.0, limma v.3.58.1 and DESeq2 v.1.40.2

## Ethics and Inclusion statements

The research is led by local researchers and include local contributions throughout all research process including the study design, study implementation, data ownership, intellectual properties and authorship for publication.

## Author contributions

RS, CChe and NC conceived, designed the study and write the original draft. NC and CChe administrated and supervised the project. NC acquired funding to collect isolates, while CChe acquired funding for sequencing and downstream analysis. NC, RS, TEW, and NS collected and identified bacterial isolates. NC, RS and RP collected clinical data. CChe, NRT, NC, RS, JT, EB and WC performed whole-genome sequencing. NPJD and NC contributed reagents. NRT and JP contributed software tools. RS, CCho, and CChe performed bioinformatics analyses. CChe, NC, NRT and JP interpret the analyses. CChe wrote the revised draft. All authors read and approved the manuscript.

