## Supplementary Information for "Genetic diversity, determinants, and dissemination of *Burkholderia pseudomallei* lineages implicated in melioidosis in northeast Thailand"

**This document provides information for Supplementary figure 1-8.**

**Supplementary table 1-4 are provided as separate files in excel.**

### Supplementary Figures

### Supplementary Figure 1

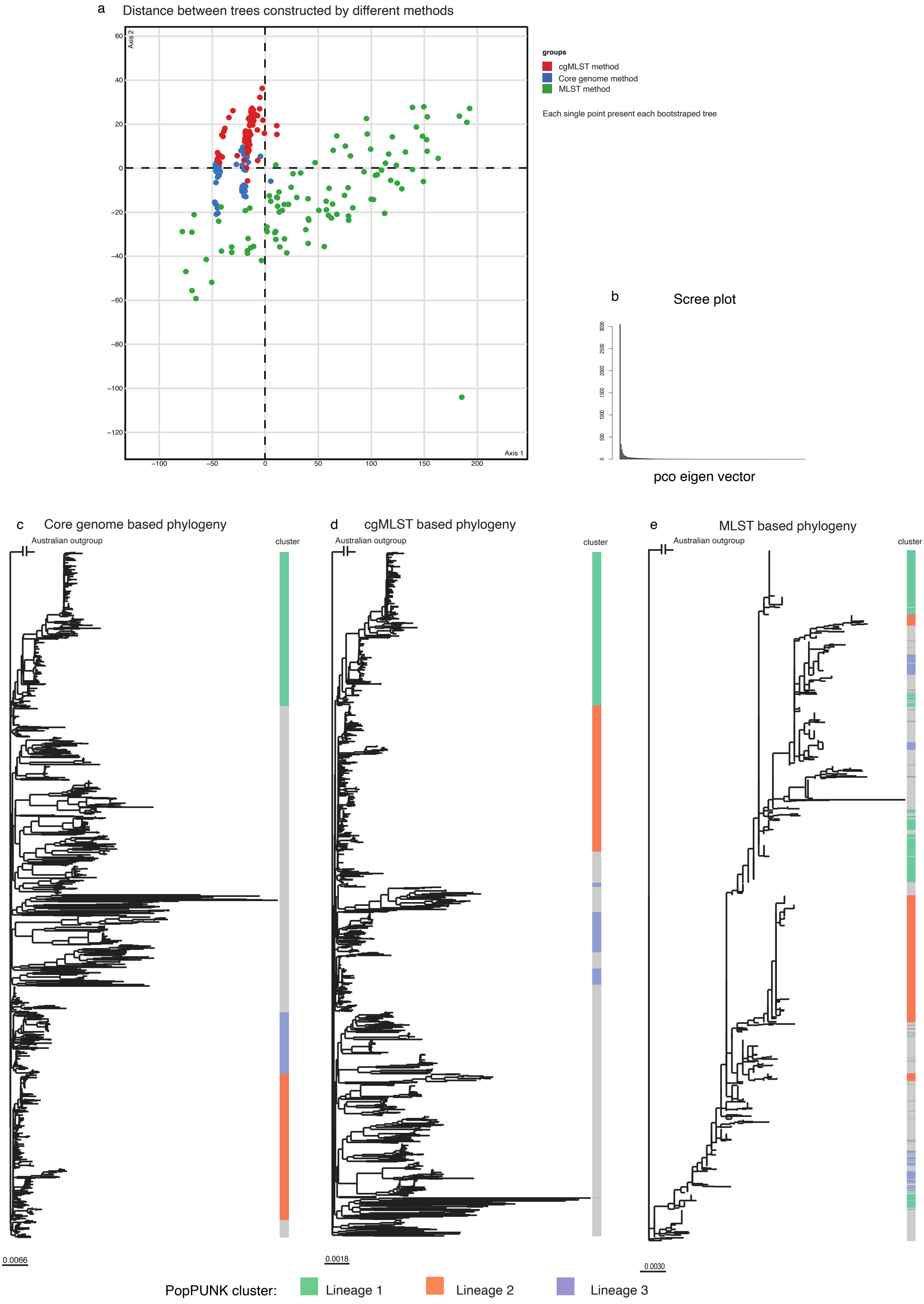

**Supplementary Figure 1. Comparison of approaches used in in outlining the population structure** (a) A scatter plot displays the first two principal components (PCs) derived from pairwise distances between trees. Each dot represents a bootstrap tree and is colour-coded by the method used to generate the tree: core genome SNP (blue), cgMLST (red) and MLST (green). (b) A scree plot summarises eigenvalues computed for each PCs. (c to d) The median phylogenetic trees constructed from core genome SNP, cgMLST, and MLST and their consistency with PopPUNK clustering method.

### Supplementary Figure 2

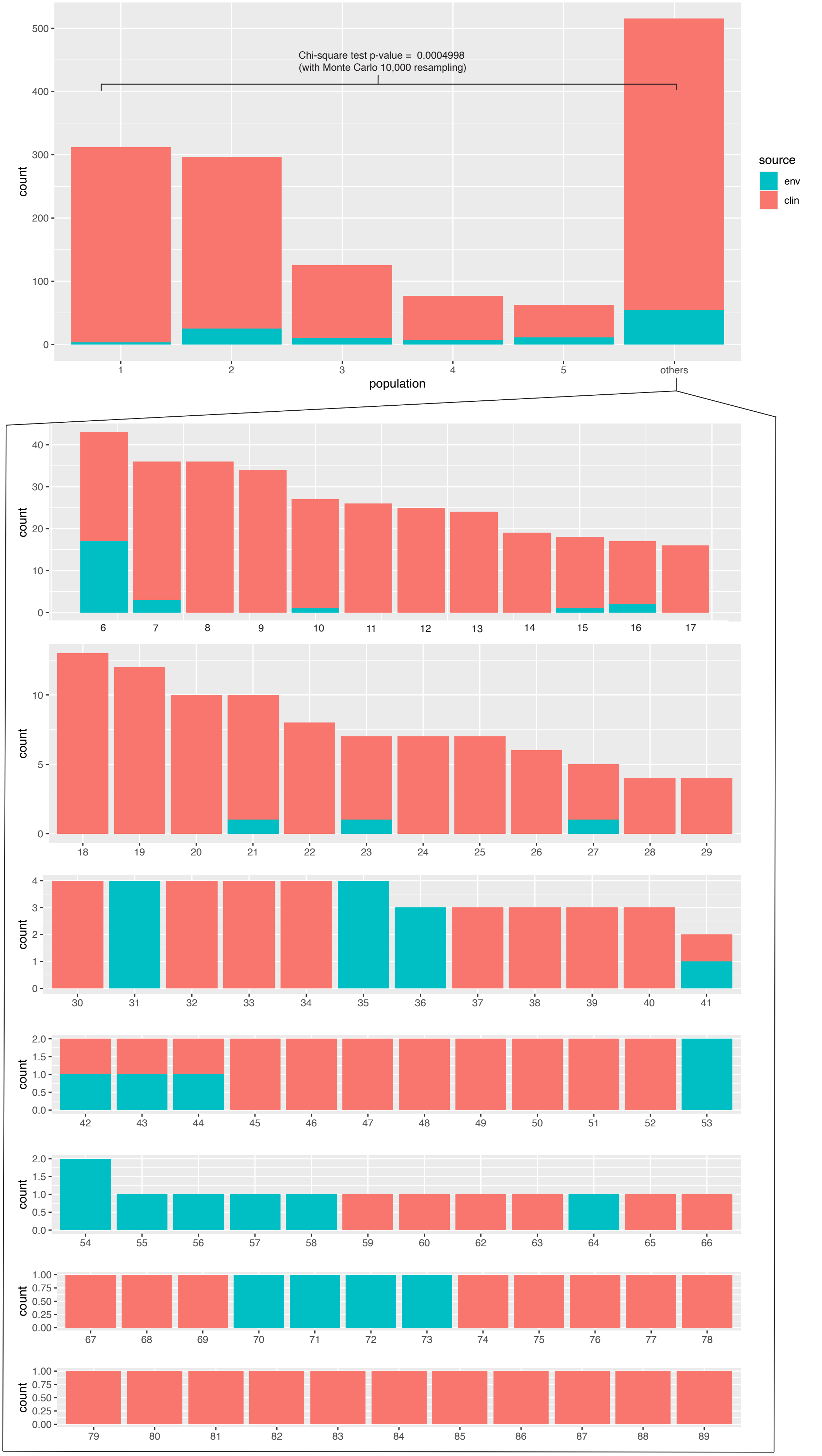

**Supplementary Figure 2. Distribution of environmental and clinical isolates by each lineage.** Barplots highlight the co-detection of environmental (green) and clinical isolates (red) across dominant lineages 1, 2, and 3.

**Supplementary Figure 3**

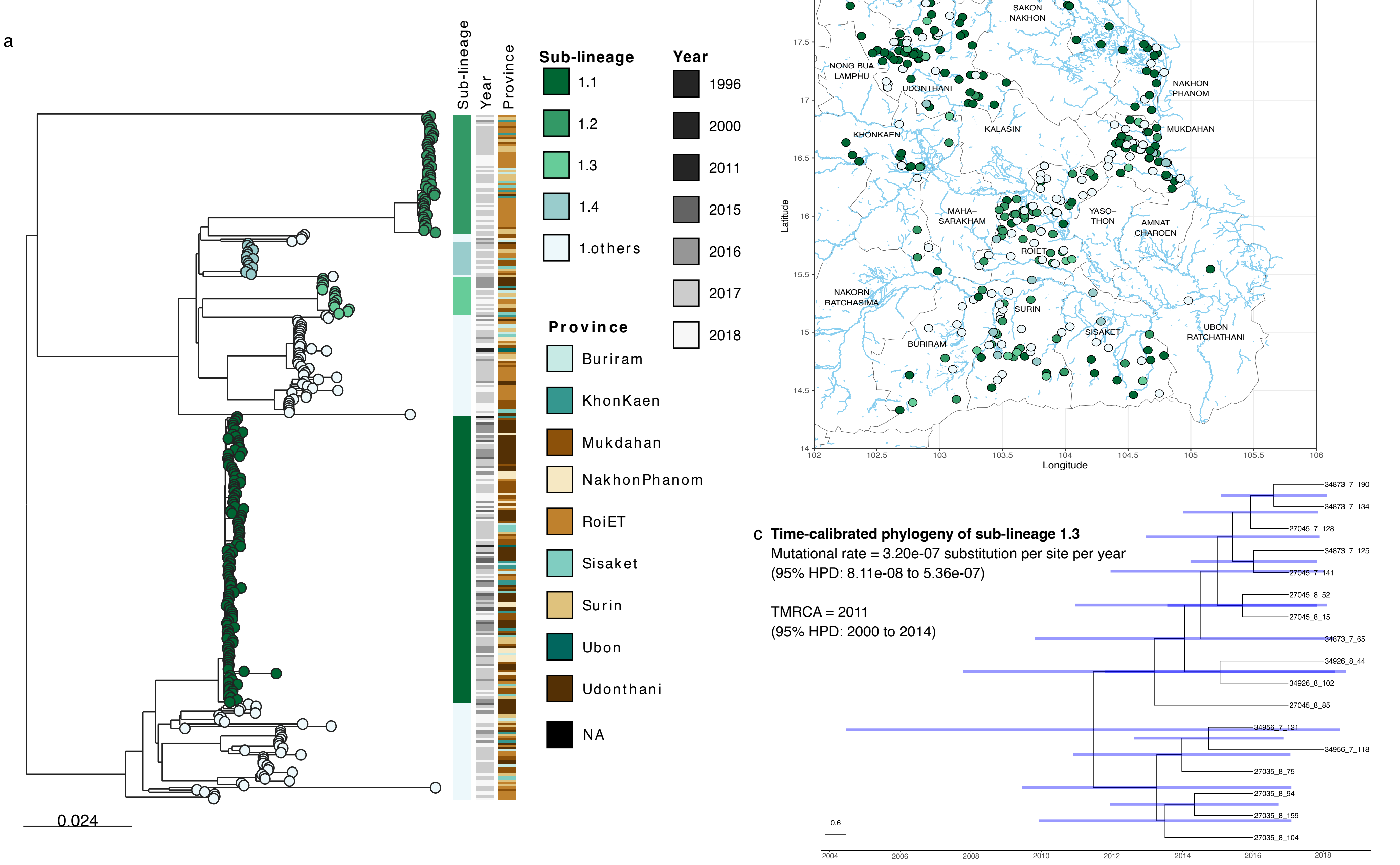


**Supplementary Figure 3. Lineage 1 specific analysis** (a) A recombination removed lineage 1 phylogeny with colour stripes displaying its sub-lineage structure, year of collection, and sampling province (left to right). (b) A map of northeast Thailand showing the distribution of each isolate and the region's river system. (c) Time-calibrated phylogeny of sub-lineage 1.3 with blue error bars indicating 95% highest posterior density interval, with the estimated mutational rate consistent with previous study.

**Supplementary Figure 4**

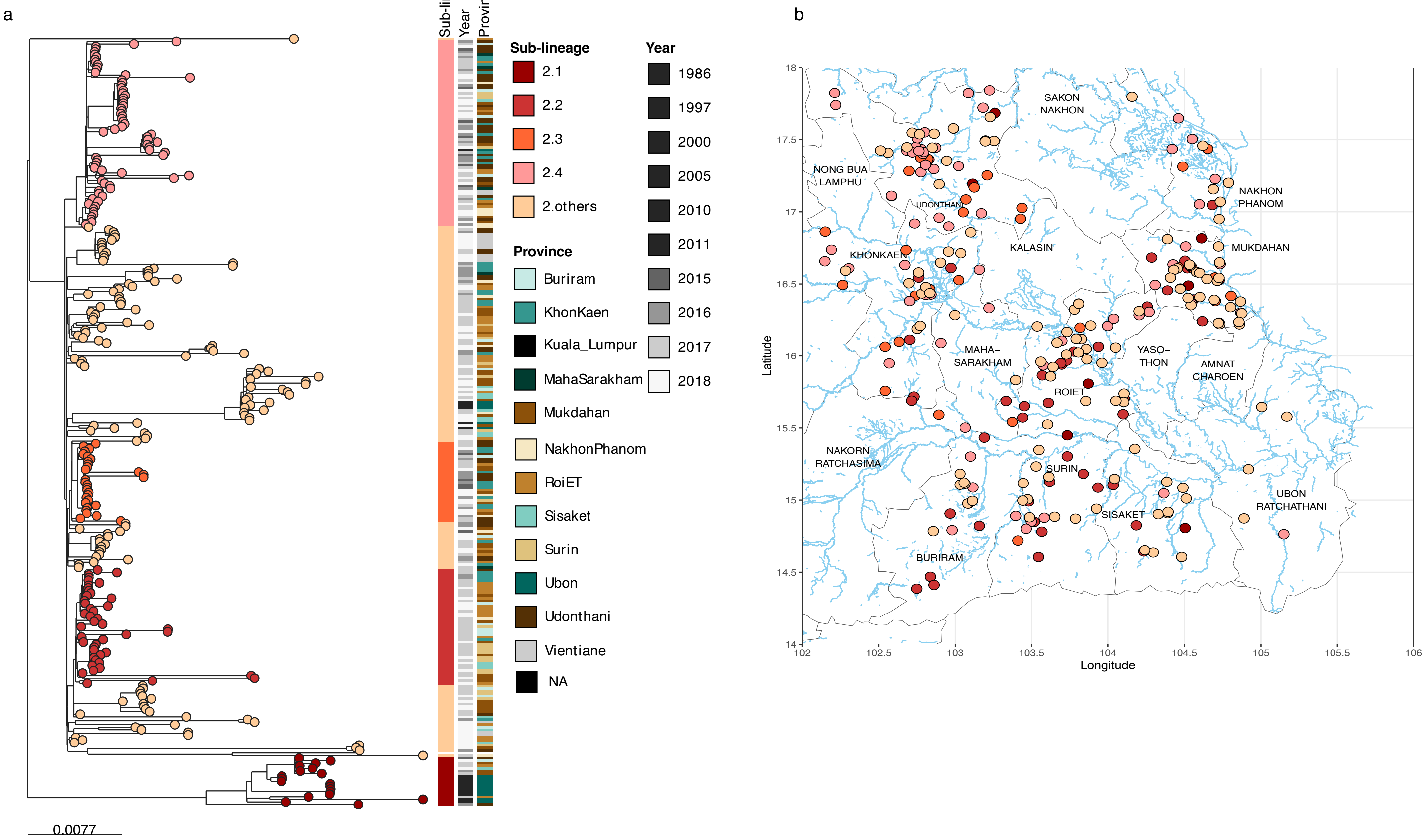


**Supplementary Figure 4. Lineage 2 specific analysis** (a) A recombination removed lineage 2 phylogeny with colour stripes highlighting sub-lineage structure, year of collection, and sampling province (left to right). (b) A map of northeast Thailand with the region's river system. Dots present the distribution of individual samples.

Supplementary Figure 5

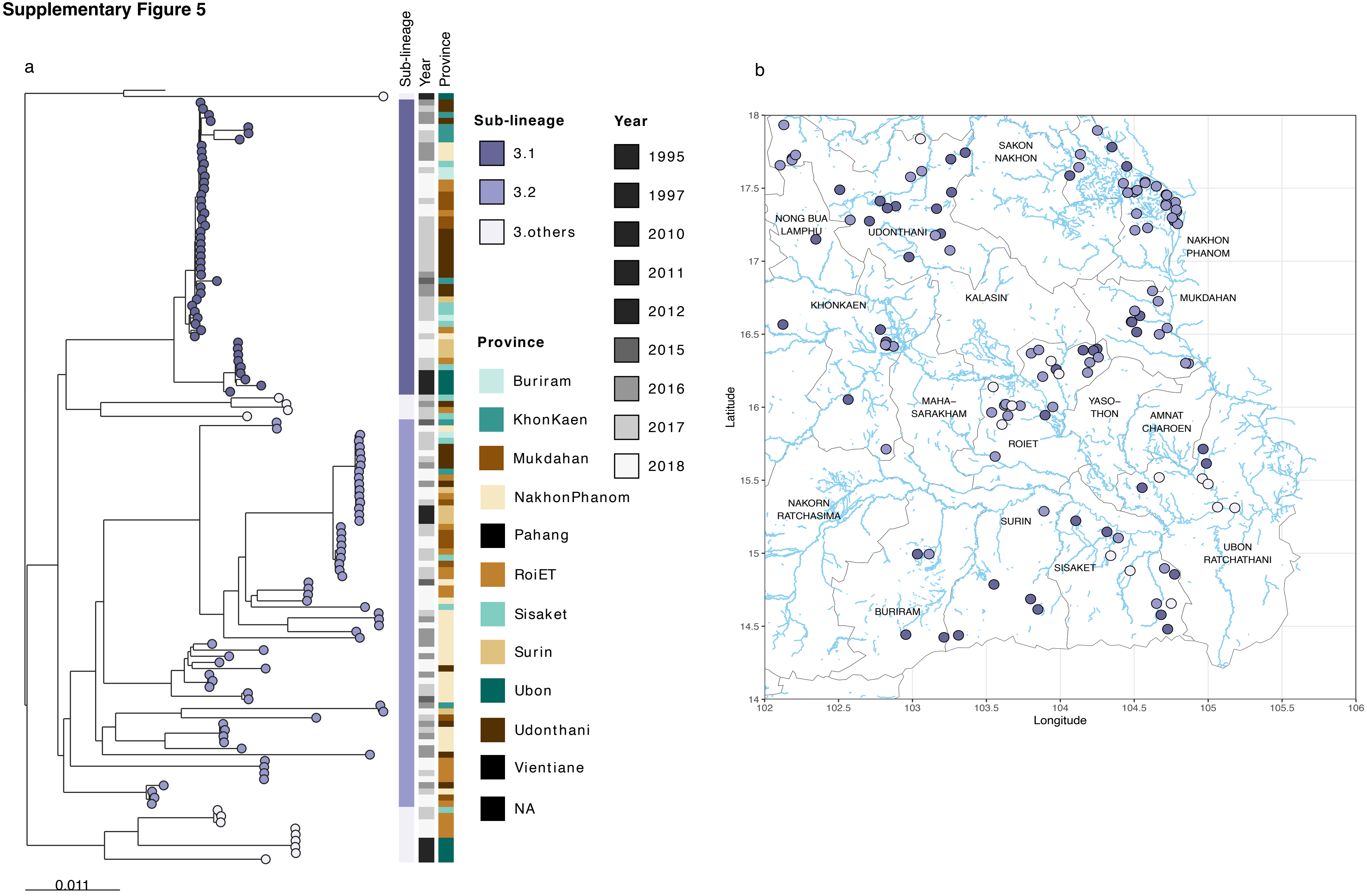


**Supplementary Figure 5. Lineage 3 specific analysis** (a) A recombination removed lineage 3 phylogeny with colour stripes highlighting sub-lineage structure, year of collection, and sampling province (left to right). (b) A map of northeast Thailand showing the distribution of each isolate and the region's river system.

**Supplementary Figure 6**

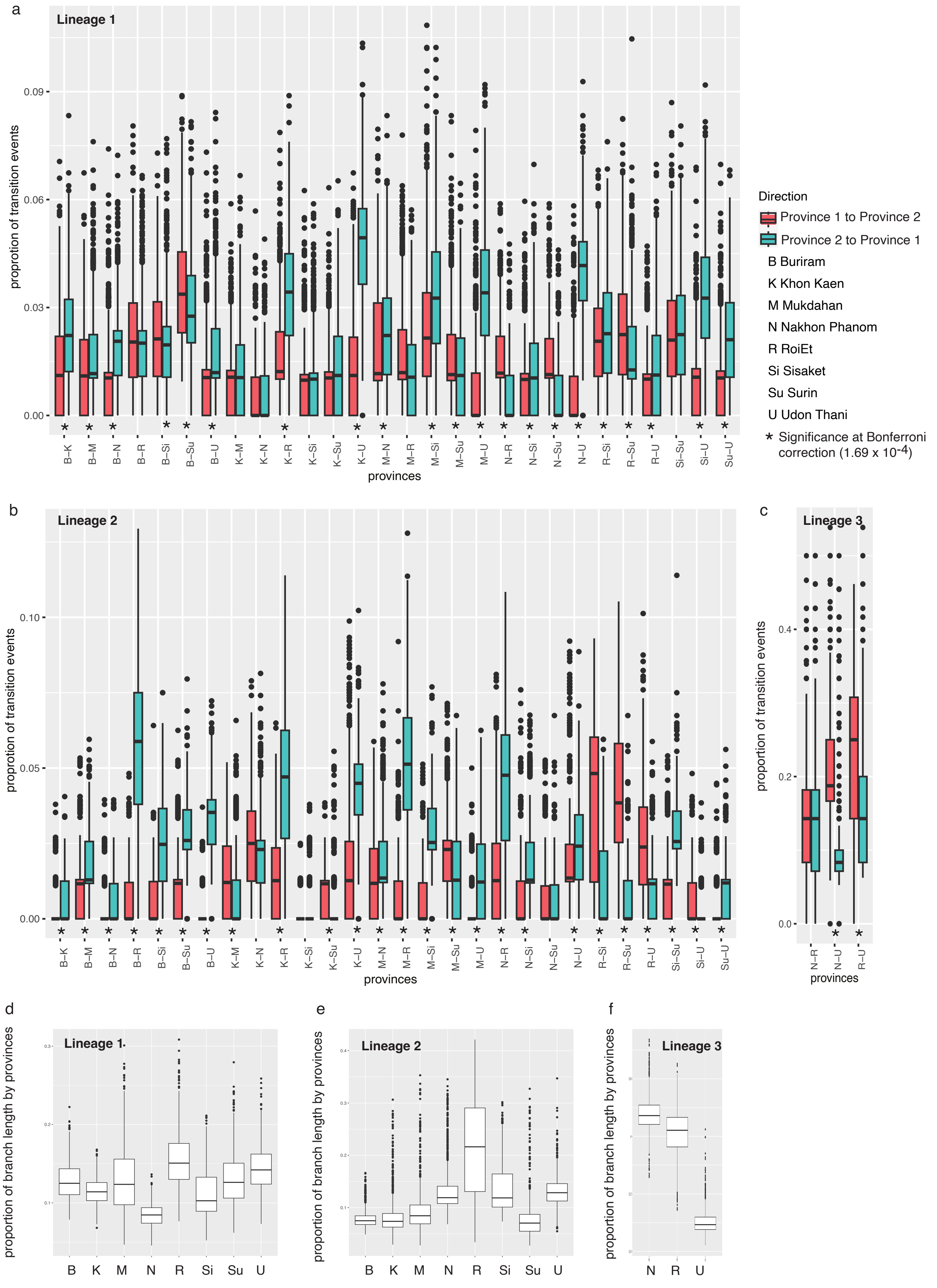

**Supplementary Figure 6. Transmission patterns and evolutionary time spent at each province**. (a to c) Proportion of transition events (Markov jumps) among provincial pairs for lineage 1, 2, and 3 respectively. The pairs were denoted as province1 – province 2, with transitions from province 1 to province 2 shown in red, and transitions from province 2 to province 1 in green. The Man-Whitney U test was conducted for each pair to assess differences in transition frequency by direction, with Bonferroni correction applied for multiple tests. (d to f) Total branch length from provincial trait reconstruction (Markov rewards) for lineage 1, 2, and 3, respectively.

**Supplementary Figure 7**

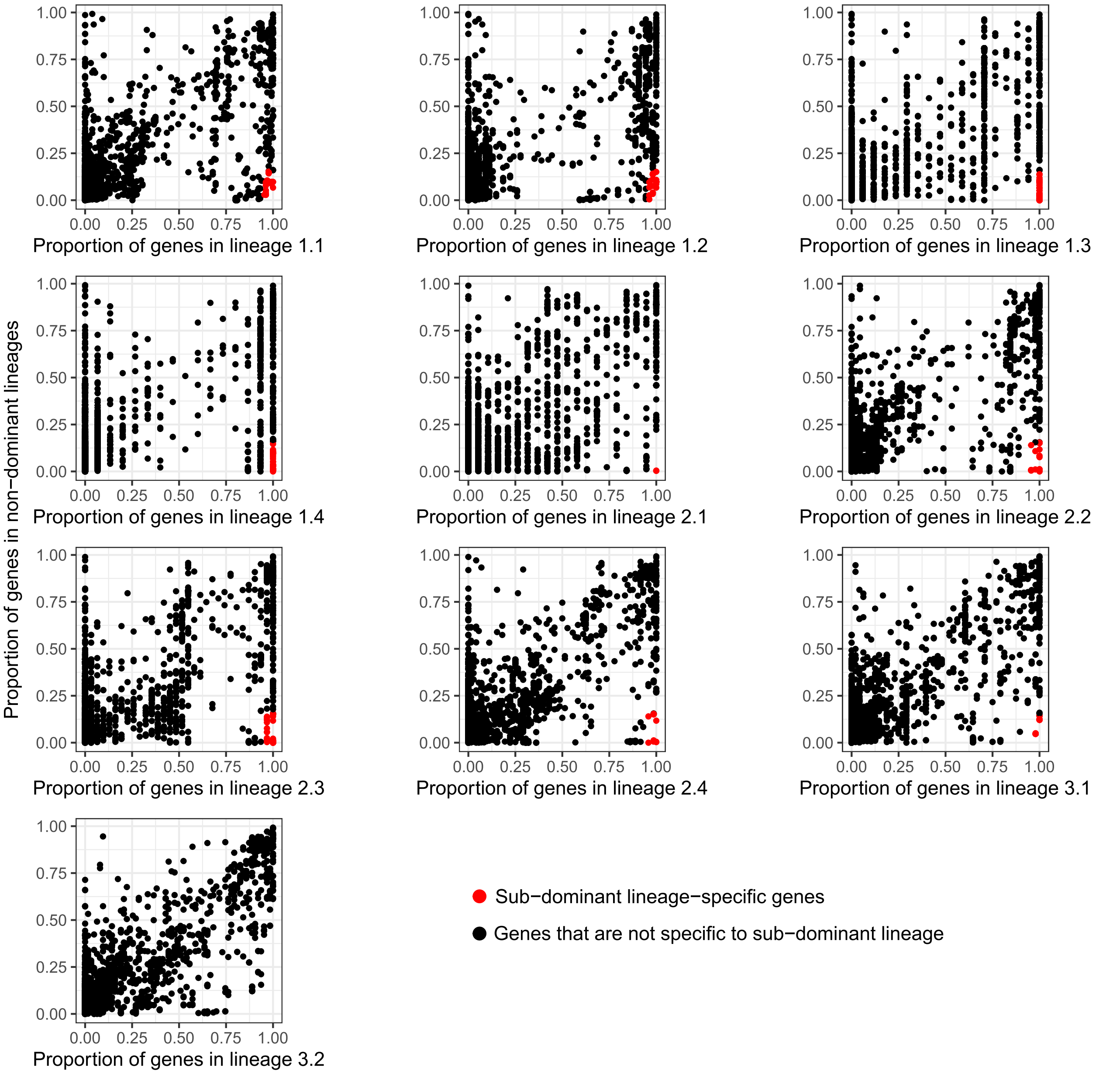


**Supplementary Figure 7. Selection criteria for lineage-specific genes**. Scatter plots show the frequency distribution of lineage-specific (red) against other genes (black), based on their distribution within the dominant lineages and their sub-lineages (horizontal axis) compared to their distribution in non-dominant lineages.

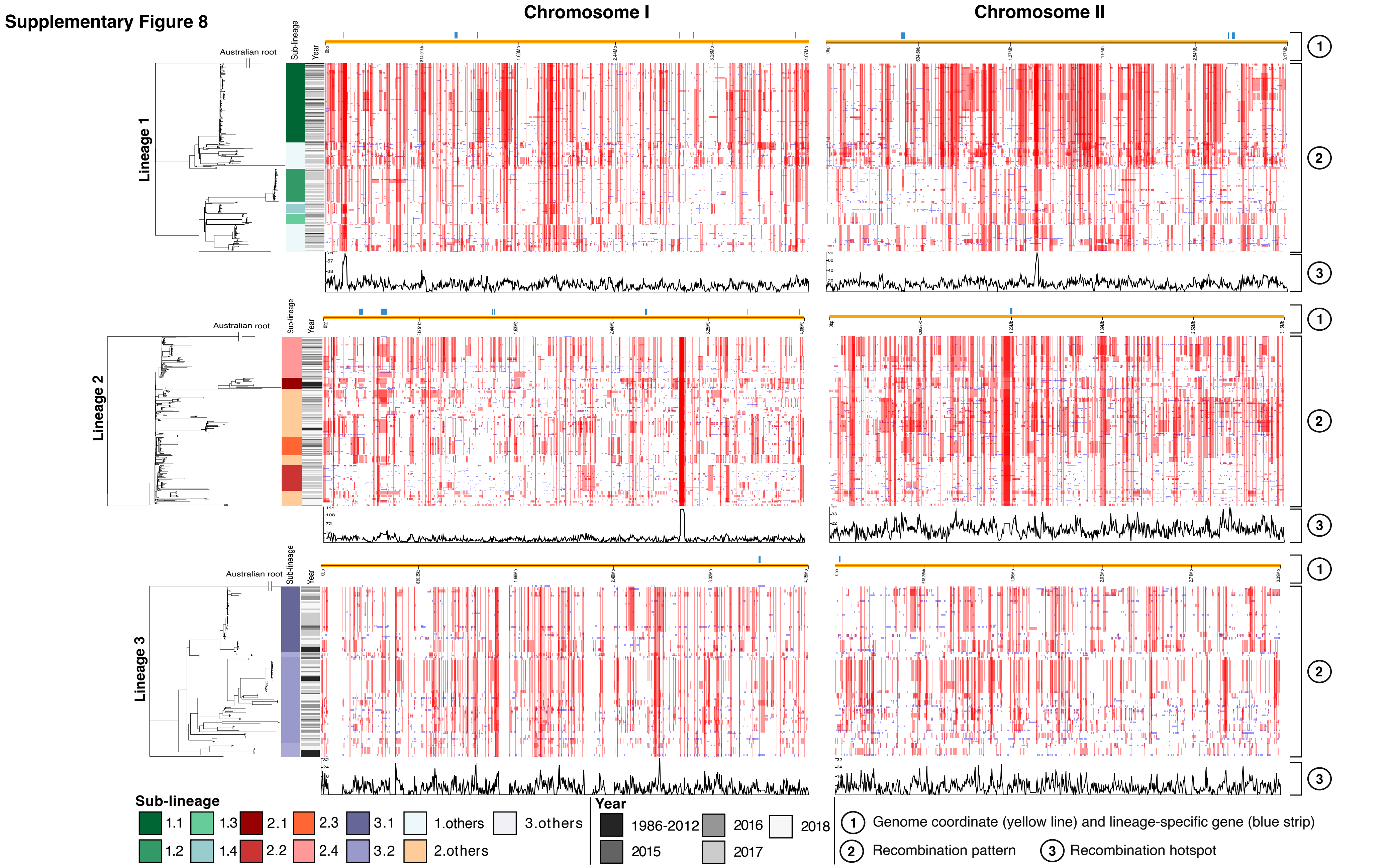


**Supplementary Figure 8. Recombination patterns detected in lineage 1, 2, and 3**. From left to right: the recombination-removed phylogeny of each lineage, a stripe representing the sub-lineage classification and sampling year, and heatmaps displaying recombination patterns identified in chromosome 1 and 2. The top orange lines mark the genome coordinates. For each lineage, their respective lineage-specific genes are highlighted in blue at the top of the panel. Each heatmap represents recombination blocks aligned with the phylogeny. Recombination events occurring at the internal nodes are coloured in red, while those occurring at the external branches are coloured in blue. The recombination hotspot is plotted at the bottom of each heatmap.
